# siRNA/mRNA payload of lipid nanoparticles quantified using fluorescence microscopy and correlation spectroscopy

**DOI:** 10.64898/2026.11.05.763588

**Authors:** Bernhard R. Kirchmair, Nikolay Stoyanov, Daria Maksutova, Thomas-Otavio Peulen, Judith A. Hagemeier, Thomas Kellerer, Thorben Cordes, Marcus Fändrich, Joachim O. Rädler

## Abstract

Lipid nanoparticles (LNPs) have emerged as the most promising vectors to deliver nucleic acids to mammalian cells. Advanced delivery strategies involving multiple nucleic acid species require a reliable determination of RNA payload and LNP size. Here, we use total internal reflection fluorescence (TIRF) microscopy and fluorescence correlation spectroscopy (FCS) to determine mRNA payload and siRNA/mRNA codelivered payload. Particularly, FCS proves to be a reliable method for payload determination, independently of fluorophore properties that can be altered during encapsulation. By varying the PEG-lipid fraction relative to the reference ONPATTRO formulation, we find that mRNA payload of enhanced green fluorescent protein (eGFP) coding mRNA increases from ∼10 with LNP size according to a power law with an exponent 1.3. Cryogenic electron microscopy (Cryo-EM) reveals that in this case blebs occur more frequently in small LNPs. For siRNA/mRNA coencapsulation the siRNA fraction within LNPs exceeds that in the initial RNA mixture, suggesting that coencapsulation is competitive and constrained by nucleic acid diffusion. Together, our findings highlight the importance of particle size and encapsulation kinetics in controlling the loading and composition of RNA payloads in LNPs.

## Introduction

Lipid nanoparticles (LNPs) are clinically established carriers for nucleic acid therapeutics. Their core– shell architecture, defined lipid composition, and ionizable lipids enable controlled particle size and near-quantitative nucleic acid encapsulation. LNPs have been optimized for highly effective mRNA delivery^1^ and were clinically validated for the first time by the siRNA drug patisiran (ONPATTRO).^2^ Since then the ON-PATTRO formulation has served as a reference point in LNP development. Many subsequent studies used lipid molar ratios related to the original ONPATTRO formulation leading to increasingly better understanding of the role of ionizable lipid and LNP physico-chemical characterization.^3,4^ Structural characterization such as small-angle X-ray scattering (SAXS)^5–8^ and cryogenic transmission electron microscopy (Cryo-EM) have revealed that the internal organization of LNPs is intrinsically coupled to how much and how tightly nucleic acid is packaged within each particle. However, the absolute number of nucleic acid molecules per LNP remains poorly quantified. Yet, understanding payload is essential, as it directly influences delivery efficiency, biological activity, and therapeutic performance.^9^ Beyond their clinical use, LNPs serve as gene delivery model systems for studying fundamental gene regulatory mechanisms. Single-cell approaches, such as live-cell imaging on single-cell arrays (LISCA), have proven particularly valuable in this context, enabling quantitative modeling of reporter gene expression following LNP-mediated mRNA delivery.^10^ These studies link payload quantity to functional protein outcome by numbers. Quantifying payload precisely is therefore critical not only for mechanistic understanding of LNP-mediated delivery but also for ensuring reproducibility, potency, and quality control in the development of LNP-based therapeutics.

Many groups have attempted to quantify the number of RNA molecules per LNP, most commonly using the ONPATTRO-type formulation (MC3/DSPC/cholesterol/PEG-lipid, 50:10:38.5:1.5) as a reference composition. Ensemble-level estimates of payload have relied on small-angle scattering (SAXS/SANS) combined with volumetric modeling of particle size, lipid composition, and encapsulated nucleic acid content^5,11,12^ or on related mass-based/molecular-volume approaches.^13,14^ For the ONPATTRO formulation specifically, these methods report payloads ranging from 5–200 mRNA copies per LNP depending on particle size,^5^ to∼30 copies,^11^∼13 copies,^12^ and ∼26 copies.^13^

Single-particle fluorescence techniques can provide more direct, particle-resolved payload measurements; methods include structured illumination microscopy,^15^ convex lens-induced confinement (CLiC) fluorescence imaging,^16,17^ multi-color cylindrical illumination confocal spectroscopy,^18,19^ and asymmetric flow field-flow fractionation coupled to SAXS and nano-flow cytometry.^20^ For the ONPATTRO formulation, single-particle approaches show low copy numbers of ∼5 luciferase-mRNA copies per LNP^15^ and ∼2.8 copies yet additionally ∼80 % empty particles,^18^ while a slight increase of 1.3–2.7 mRNA copies per LNP with 20 % to 35 % empty particles is seen for distinct formulation buffers.^17^ For siRNA much higher copy numbers are reported. Interestingly, a drop from ∼100 to ∼10 siRNA copies per LNP is found as PEG-lipid content increases from 0.5 mol % to 4.5 mol %.^19^

All these findings underscore the substantial variance in the determined payload, next to the fact that many particles are only partially loaded, despite seemingly uniform size distributions. Recently, the reported mRNA payloads were converted into a volume-normalized effective nucleotide density leading to a significant harmonization.^21^ Additionally, Kamanzi et al.^22^ used simultaneous size, lipid, and cargo fluorescence imaging (CLiC-ALEX) across multiple clinically relevant formulations to show that the mRNA payload heterogeneity correlates with particle structure, including the prevalence of bleb morphologies. That is, bleb-positive LNPs, characterized by localized protrusions of the lipid envelope, are more likely to contain mRNA. This variability poses challenges in cell transfection experiments since the amount and stochastic variation of the delivered cargo is poorly understood. Strikingly, there has been no multi-technique approach to confirm payload independently. Also, the payloads of different RNA cargos, such as mRNA and siRNA, have not been compared and no study of payload in co-delivery within one LNP has been performed.

Here, we combine quantitative single-particle fluorescence microscopy and fluorescence correlation spectroscopy (FCS) to determine mRNA and siRNA payload in LNPs. First, we used total internal reflection fluorescence (TIRF) microscopy to compare the fluorescence brightness of individual immobilized LNPs with single RNA molecules. This provides a direct quantification of payload at the single-particle level. Simultaneously, the assay reveals important limitations arising from environment-dependent fluorophores used for mRNA visualization. Second, FCS is employed as an intensity-independent approach based on particle counting in solution, in which the reduction of the number of fluorescent particles is determined when mRNA becomes assembled into an LNP. By combining FCS with dynamic light scattering (DLS), both particle concentration and hydrodynamic size are accessible, enabling direct determination of the average number of RNA molecules per LNP. Using these methods, we find a consistent mRNA payload of 7 ±3 by TIRF and 12 ±3 by FCS and an siRNA payload of 380± 50, in LNPs formulated according to the ONPATTRO composition. Secondly, by adding PEG lipid to the formulation, we systematically increased LNP size, anticipating a volumetric, “cubic” scaling. Surprisingly, we found a scaling of payload with diameter that follows a power of 1.3. Supported by Cryo-EM, we propose blebs as explanation for this weak scaling and the decrease of the payload with the LNP size. In the case of siRNA/mRNA coencapsulation with defined molar fraction, we observed an enhanced presence of siRNA in mixed LNPs. The observed siRNA/mRNA ratios within LNPs are explained within a kinetic condensation model that assumes a tenfold higher association rate constant for siRNA than for mRNA, consistent with the approximately tenfold higher diffusion coefficient of siRNA.

## Results and discussion

### Microfluidic formulation of LNPs

Lipid nanoparticles (LNPs) were formulated using a composition based on the well-known ONPATTRO reference formulation (Figure 1). LNPs were formed by rapid mixing of a lipid ethanol phase and mRNA-loaded aqueous phase in a ratio of 1:2 (for further details see Materials and methods). Enhanced green fluorescent protein (eGFP) coding mRNA (996 nt) that had been labeled with Cy5 was used as nucleic acid cargo. RNA encapsulation efficiency exceeded 90 % for all preparations, as quantified by the RiboGreen assay.

**Figure 1:**
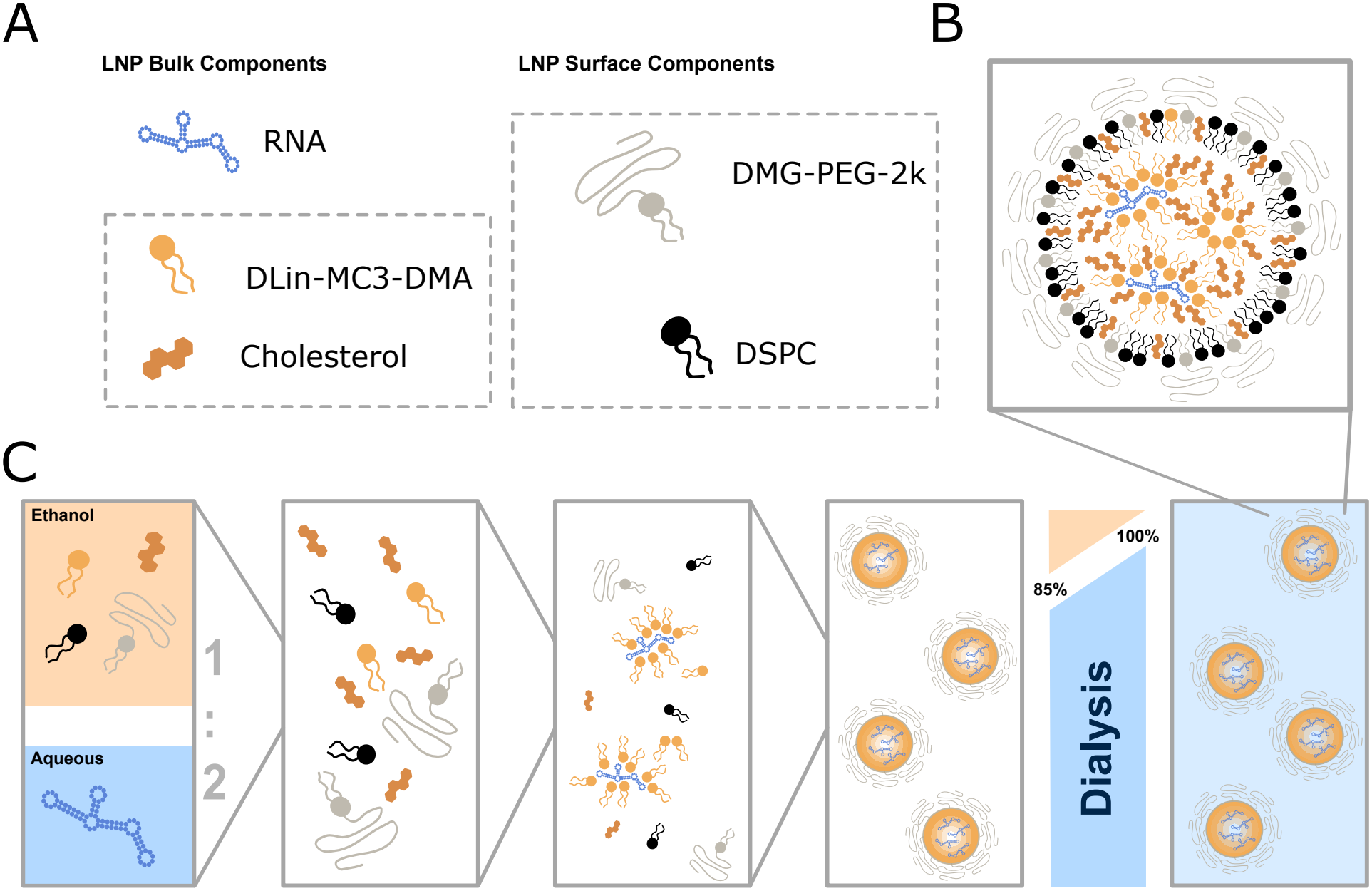
Microfluidic formulation and self-assembly of LNPs. (A) Aqueous RNA solution and ethanol-based lipid solution are rapidly mixed in a microfluidic device. (B) Initial isotropic distribution of RNA and lipids in the mixed water:ethanol (2:1) solvent. (C) Condensation of RNA and ionizable lipid via electrostatic interactions. The self-assembly of PEG-lipid coated LNPs is driven by hydrophobic forces. In the final step remaining ethanol is extracted by dialysis. LNPs are shown in the lower right corner as a schematic cartoon with a condensed lipid core phase (ionizable lipid, cholesterol and RNA) and a surface lipid monolayer composed of mainly the surface-active lipids (DMG-PEG-2k and helper lipid DSPC).

### Quantitative single particle fluorescence microscopy

Quantitative single-particle fluorescence microscopy was used to determine the mRNA payload per lipid nanoparticle by comparing fluorescence intensities of individual LNPs and single mRNA strands. Although the physical sizes of both species are below the diffraction-limited resolution of an optical microscope, the number of fluorophores per diffractionlimited spot is encoded in the measured brightness, enabling a relative quantification. To this end LNPs were immobilized using biotinylated PEG and a neutravidin-coated glass slide (Figure 2A). As a reference, bare single mRNA molecules were hybridized to a biotinylated polyT strand. Surface-tethered Cy5-labeled mRNA and LNPs were then imaged under TIRF illumination showing randomly distributed fluorescent spots (Figure 2B,C). LNPs appeared as bright spots with heterogeneous intensity, while single mRNA molecules were dimmer, yet with a more homogeneous intensity distribution. Analysis of localization data in Thunder-STORM yielded brightness histograms for LNPs and mRNAs (Figure 2D). Here, the intensities of the spots were integrated and plotted as particle fluorescence intensity distributions. It has to be noted that the left part of the histograms is influenced by the choice of threshold values used for emitter identification. Under similar thresholding conditions, the LNP intensity distribution was broad, yet featured a clear maximum value under the given thresholding conditions. The bare mRNA molecules, due to their lower brightness, showed a distribution with a steep rise, i.e., brightness values just above background, and an exponential decay indicating single-molecule characteristics featuring a discrete number of Cy5 emitters (Figure 2D).

**Figure 2:**
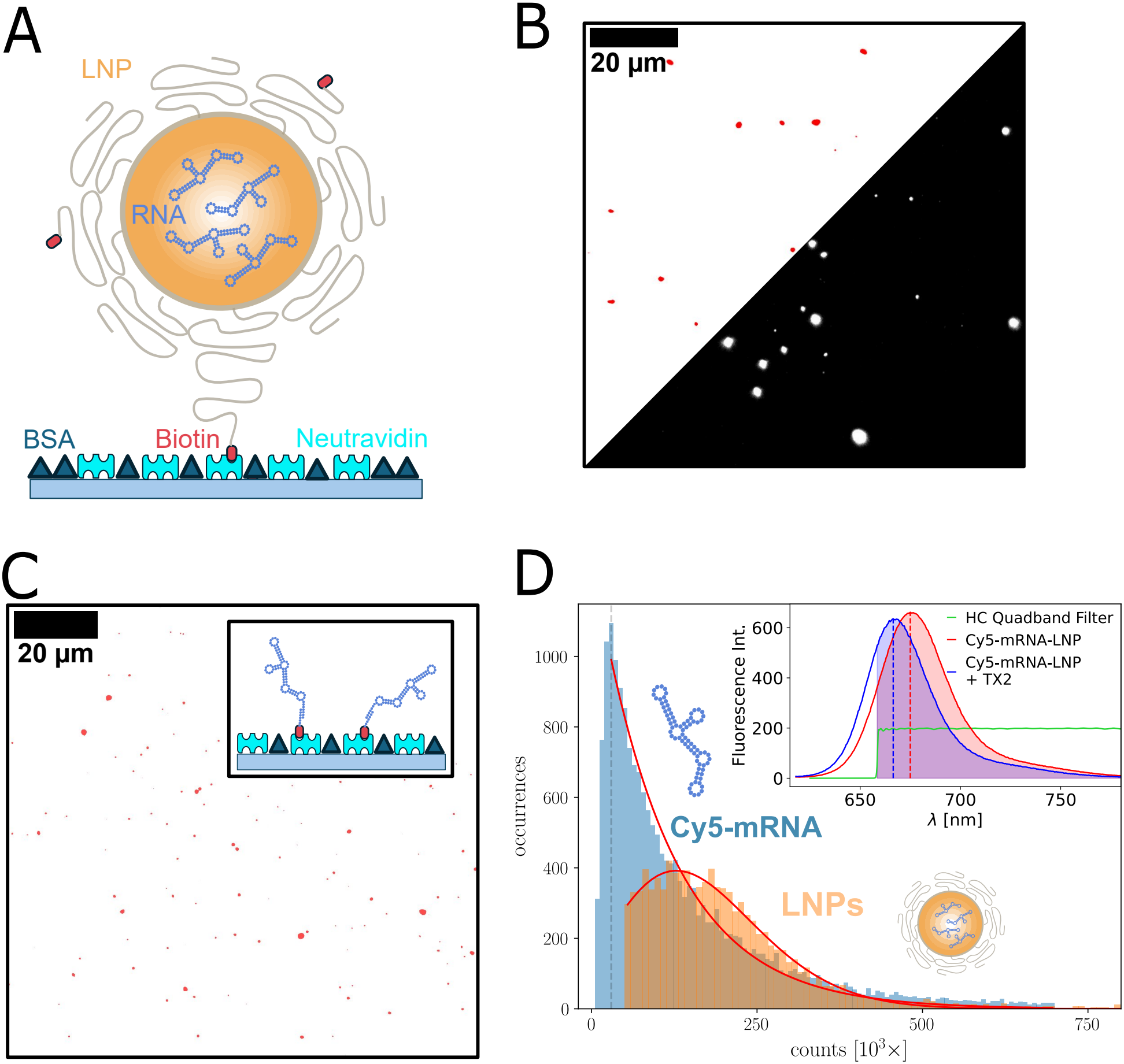
Number of mRNAs per LNP determined by quantitative fluorescence imaging. (A) Cartoon showing LNPs anchored to a BSA-coated glass slide via biotin–neutravidin linkers. (B) Fluorescence micrograph of surface-anchored Cy5-mRNA-LNPs. (C) Inverted intensity micrograph of biotin-polyT-anchored mRNAs on a neutravidin-coated glass slide. Inset: cartoon showing mRNA–polyT linkage. (D) Intensity histogram of LNPs shows an increased brightness compared to bare mRNA. Exponential mRNA fit and Poissonian LNP fit shown in red. Inset: solvatochromic shift of Cy5-mRNA inside the LNP and after Triton-induced release of mRNA from the nanoparticles.

The single-molecule brightness ⟨*I*(mRNA = 1) ⟩ is determined from the maximum of the intensity distribution using a Gaussian kernel density estimator. The LNP distribution suggests stochastic loading of the particles, in which Poissonian fitting gave a mean LNP brightness *I*(LNP) . Since LNPs contain a large number of emitters, the brightness is lifted well above the background values giving rise to the Poisson distribution.

Importantly, we noted that the fluorophore photophysics, i.e., both spectra and fluorescence lifetime, were altered upon encapsulation since both the chemical environment and viscosity of the LNP interior strongly differ from water (Figure S1). We found that Cy5 exhibits a hypsochromic shift of (8.5 ±0.5) nm upon encapsulation into LNPs, obtained by comparison of the emission spectra before and after Triton X-100-induced disruption of LNPs at constant mRNA concentration (Figure 2D inset and Figure S1). We observed a subtle reduction in the total emission intensity of the dye after addition of TX100 to LNPs (Figure S1). In addition, LNP-encapsulated Cy5 showed a higher fluorescence lifetime relative to Cy5 on free mRNA (after addition of 2 % TX100), consistent with a reduced non-radiative decay (cis–trans isomerization) due to restricted conformational flexibility or increased viscosity.^23,24^ This was qualitatively in line with the reduced intensity of Cy5 after addition of TX100, yet the magnitude of lifetime and intensity changes were not identical, indicating that the brightness comparison of mRNA-Cy5 in free and LNP-bound form is intricate. The spectral shift also affects detection due to the finite transmission window of the HC quadband filter used in TIRF microscopy. As a result, the detected intensity of LNP-encapsulated Cy5 is increased relative to free Cy5. This effect was accounted for by introducing an optical transmission factor *ψ*_T_ = 1.3, obtained from the ratio of integrals of emission spectra weighted by the filter transmission function. Hence the average mRNA payload per LNP based on fluorescence intensity including spectral shift correction is given by Eq. 1.

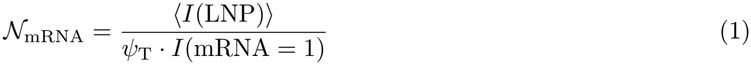

Taking ⟨*I*(LNP)⟩ from the Poisson fit in Figure 2D and *I*(mRNA = 1) from the exponential distributions as explained above, we obtain N_mRNA_ = 7 ± 3, a value close to other literature values.

### Determination of RNA payload using FCS

Since mRNA payload measurement based on LNP brightness required diligent analysis of the fluorescence shift in the LNP moiety, we validated the results using a complementary, intensity-independent approach. Fluorescence correlation spectroscopy (FCS) determines LNP size and particle concentrations from intensity fluctuations in a well-defined confocal detection volume (Figure 3A).^25^ The average effective particle number in the detection volume is given by the inverse of the zero-lag autocorrelation amplitude, ⟨ *N*⟩ = 1*/G*(0) (Figure 3B).^26^ For determination of RNA payload, the particle numbers of free mRNA before complexation and particle numbers of mRNA-loaded LNPs after complexation are compared. The decrease of the 1*/G*(0) value indicates how many mRNAs were encapsulated per LNP. Identical mRNA concentration of the two samples is ensured by measuring the absorption at 260 nm with a NanoDrop spectrophotometer. The average number of mRNA molecules per LNP is then given by the particle number quotient of free mRNA, ⟨ *N*_mRNA_ ⟩, and mRNA complexed in LNPs, ⟨ *N*_LNP_ ⟩. To verify that this relation is independent of concentration and instrument dynamic range, measurements were performed over a dilution series (Figure 3C). Free mRNA and mRNA-loaded LNPs were diluted over a concentration range of *c* = 10^*−*6^ to 10^*−*1^ mg mL^*−*1^ mRNA. As seen in Figure 3C the particle number density is proportional to mRNA concentration, both for mRNA, 1*/G*_mRNA_(0) = *A*_mRNA_ *c*, and LNPs, 1*/G*_LNP_(0) = *A*_LNP_ *c*. The ratio between the two proportionality coefficients provides the mRNA payload.

**Figure 3:**
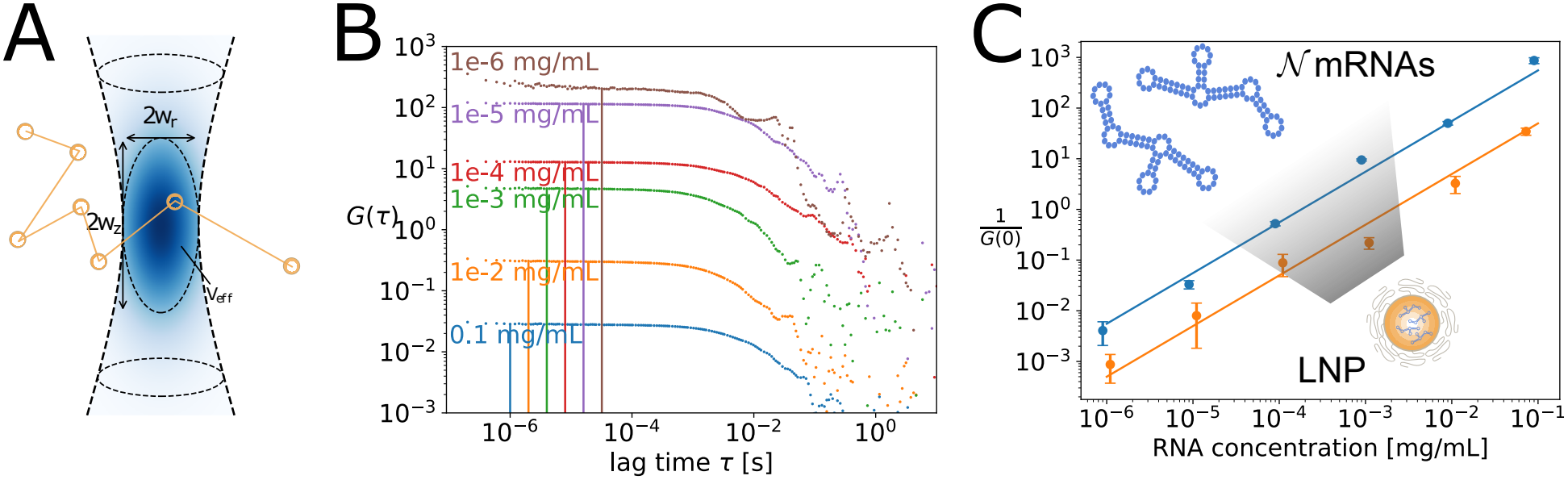
mRNA payload per LNP determined by FCS. (A) Schematic view of fluorescent particles passing through the confocal volume. (B) Fluorescence autocorrelation function of mRNA LNPs at concentrations ranging from 10^*−*1^ to 10^*−*6^ mg mL^*−*1^. The amplitude is inversely proportional to the concentration of the nanoparticles or mRNA. (C) Inverse autocorrelation function amplitudes 1*/G*(0) as a function of free mRNA (blue) and LNP (orange) concentrations in a double-logarithmic plot, each with linear fit of slope 1 (*N* = 3).

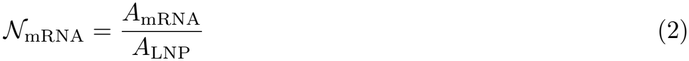

We determined the amplitudes via a linear fit in the log–log representation (Figure 3C). Using this approach, we obtain an average payload of *N* _mRNA_ = 12 ±3 mRNA molecules per LNP for the standard ONPATTRO formulation. This value lies slightly above the result from the quantitative TIRF approach, though both remain within overlapping error bars. We expect the FCS-based result to be more robust, as the normalized autocorrelation value *G*(0) reflects purely statistical fluctuations at the single-particle level, making it in principle independent of fluorophore brightness or photophysical properties in the limit of narrow intensity distributions. Batch-to-batch variations in the mRNA labeling quality may also contribute to the observed discrepancy, but overall, the agreement between the two methods strengthens confidence in the determined payload of mRNA per LNP.

### Tuning LNP size by compositional variation

Having established the mRNA copy number per LNP for a reference formulation, we next examine to what extent the nucleic acid payload can be modulated. One straightforward strategy is to vary particle size, as larger LNPs may accommodate higher RNA copy numbers. LNP size has previously been shown to influence mRNA vaccine immunogenicity for diameters between 50 nm to 150 nm,^27^ an effect commonly attributed to changes in transfection efficiency rather than to differences in mRNA content. FCS was used to determine the hydrodynamic radius *R*_*H*_ of LNPs from their characteristic diffusion times and the Stokes relation (see Materials and methods and Figure 4A).

**Figure 4:**
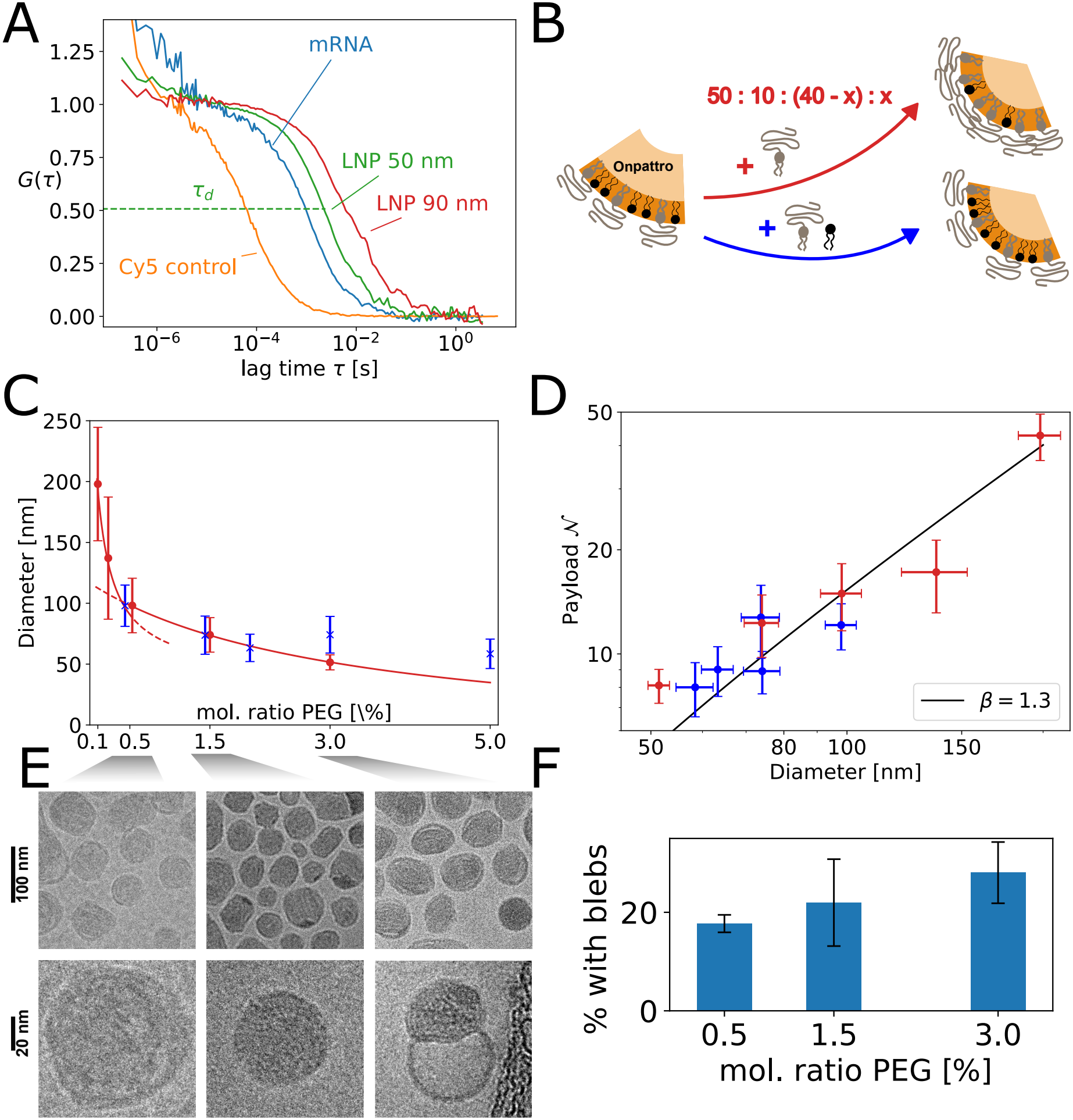
Tuning LNP size and mRNA payload by variation of LNP lipid composition. (A) Normalized fluorescence autocorrelation functions of various LNPs showing determination of particle size by means of diffusion time *τ*_*d*_. (B) LNP size is tuned by adjusting the surface-to-volume fraction. Here we compare PEG crowding (red) and fixed surface composition formulations where ratios within surface and bulk lipids are kept constant (i.e., MC3/cholesterol = const. and PEG/DSPC = const., blue). The percentages given refer to (MC3:DSPC:Chol:PEG), where *x* is the tunable parameter (see Materials and methods). (C) LNP diameters measured via FCS as a function of PEG-lipid content. PEG crowding allows size variations from 50 nm to 180 nm diameter (*N* = 3). (D) Log–log representation of mRNA payload versus LNP diameter *d*. A linear fit in the log–log representation yields a scaling exponent of *β* = 1.3. (E) Cryo-EM images of LNPs at 0.5 %, 1.5 % and 3 % PEG, each at two magnifications. At higher molar fraction of PEG, i.e., smaller diameter, blebs are more prevalent. (F) Percentage of LNPs showing blebs evaluated from data shown in (E) for 0.5, 1.5 and 3 % PEG, respectively.

LNP size was tuned by changing the lipid composition of the formulation, which affected the ratio of surface-active lipids to the total lipid volume. As shown in the inset of Figure 4B, increasing surface components decreases the size of LNPs. This is experimentally achieved by (i) increasing the PEG lipid molar ratio at the expense of cholesterol (further referred to as “PEG crowding”) or (ii) by increasing the PEG/DSPC content, simultaneously keeping the PEG/DSPC molar ratio and hence the surface composition fixed (see Materials and methods for molar fractions).

Experimentally, (i) was found to be significantly more effective in tuning LNP size than (ii) (Figure 4C).

(i) yielded diameters from approximately 60 nm to 180 nm, beyond which particle stability was compromised (see Figure S2). Over this range, the PEG-lipid content was varied between 5 mol % and 0.1 mol %. In contrast, fixed surface composition formulations did not permit efficient size modulation, indicating that LNP curvature is primarily governed by the PEG brush.^28^

Simultaneous FCS measurements of LNP size and nucleic acid payload enabled direct investigation of their relationship. For core–shell particles with a shell of thickness *r*_0_ and a diameter of *d*, one would expect payload to scale as *N*∝ (*d*− 2*r*_0_)^3^ if RNA was distributed uniformly within the core. A shell thickness of *r*_0_ = 6 nm has been found by Yanez Arteta et al.^5^ Applying the same relation with the exponent as fitting parameter, we observe a scaling

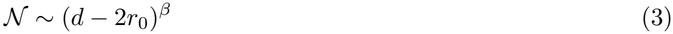

with *β* = 1.3 at a goodness of fit of *R*^2^ = 90 % (Figure 4D), suggesting that nucleic acids are not homogeneously distributed within the LNP interior. This sub-volumetric behavior is consistent with localization of RNA to internal interfaces or surface-associated substructures rather than bulk filling of the particle core.^29^ Our results are in contrast with those from single-particle analysis of siRNA LNPs by Li et al.,^19^ who used a coupled spectroscopy–chromatography method to resolve loading–size correlations at single-particle resolution. They reported a scaling factor *β* of 2.8 across different formulations for siRNA loading. At the same time they reported that within any single fixed-PEG formulation, individual-particle siRNA load-ing scaled with *β* parameters far exceeding 3 for spherical LNPs, i.e., they observed a “supervolumetric” scaling. They attributed this to a two-mode packing mechanism: below a minimum diameter (∼ 14 nm), siRNA is confined to a quadratically scaling surface layer, while larger particles can additionally accommo-date siRNA-condensed “inverted micelles” in their core. The number of the latter grows faster than particle volume, thereby exceeding cubic scaling at the single-particle level. Our ensemble FCS measurements are methodologically most comparable to their cross-formulation analysis, yet we find a considerably lower expo-nent (1.3 vs. 2.8) for mRNA than they report for siRNA. We attribute this discrepancy to the substantially larger contour length and lower flexibility of the 996 nt mRNA cargo relative to a 20 bp siRNA duplex. The lower flexibility of mRNA likely prevents efficient core condensation into compact micelle-like structures of the type described for siRNA, particularly at smaller LNP sizes. Cryo-EM imaging shows the liquid-crystalline core of LNPs (Figure 4E) with a tendency towards decreasing size with increasing PEG content(see Figure S3). Moreover, Cryo-EM reveals considerable heterogeneity with a fraction of LNPs exhibiting bleb-like structures. Evaluation of the percentage of bleb-forming LNPs shows that a higher PEG-lipid content correlates with a higher prevalence of LNP surface blebs (Figure 4F). These blebs could arise as a consequence of excess surface area: when the surface-to-volume ratio is high, the surplus PEG-lipid cannot be accommodated by a spherical shell and leads to bleb formation. Importantly, the existence of blebs provides a structural explanation why payload does not scale with particle volume, resulting in *β* parameters *<* 3. Smaller, more heavily PEGylated particles are more likely to carry blebs, a considerable addition to their core volume, and the densely packed RNA within these structures contributes to overall payload in a way that is not captured by a simple volumetric scaling argument.

### siRNA-loaded LNPs

Finally, we examined whether the nucleic acid payload per LNP can be generalized across different RNA types. Small interfering RNA (siRNA) is a double-stranded RNA molecule. In contrast to the extended structure of the 996 nt mRNA studied above, 20 nt long siRNA is akin to a 6 nm long rigid rod. Figure 5A shows FCS data obtained for Atto390-labeled siRNA free in solution and complexed in LNPs as a function of defined initial siRNA concentrations. Applying the same FCS-based analysis to siRNA-loaded LNPs yielded an average copy number of *N* _siRNA_ = 380± 50 for the ONPATTRO formulation at identical nitrogen-to-phosphate (N/P) ratio, defined as the molar ratio of ionizable lipid nitrogen atoms to nucleic acid phosphate groups (Figure 5A). The siRNA-LNP size did not significantly differ from mRNA LNPs. Due to the smaller molecular size and higher linear charge density of 20 bp siRNA compared to 996 nt mRNA, the mRNA molecule is 25 times more charged. Interestingly, expressing payload in terms of the number of nucleotides, we find (15 200± 2000) nt (siRNA) and (11 952± 3000) nt (mRNA). Hence, at equal N/P ratio a comparable number of nucleotides is encapsulated.^21^

**Figure 5:**
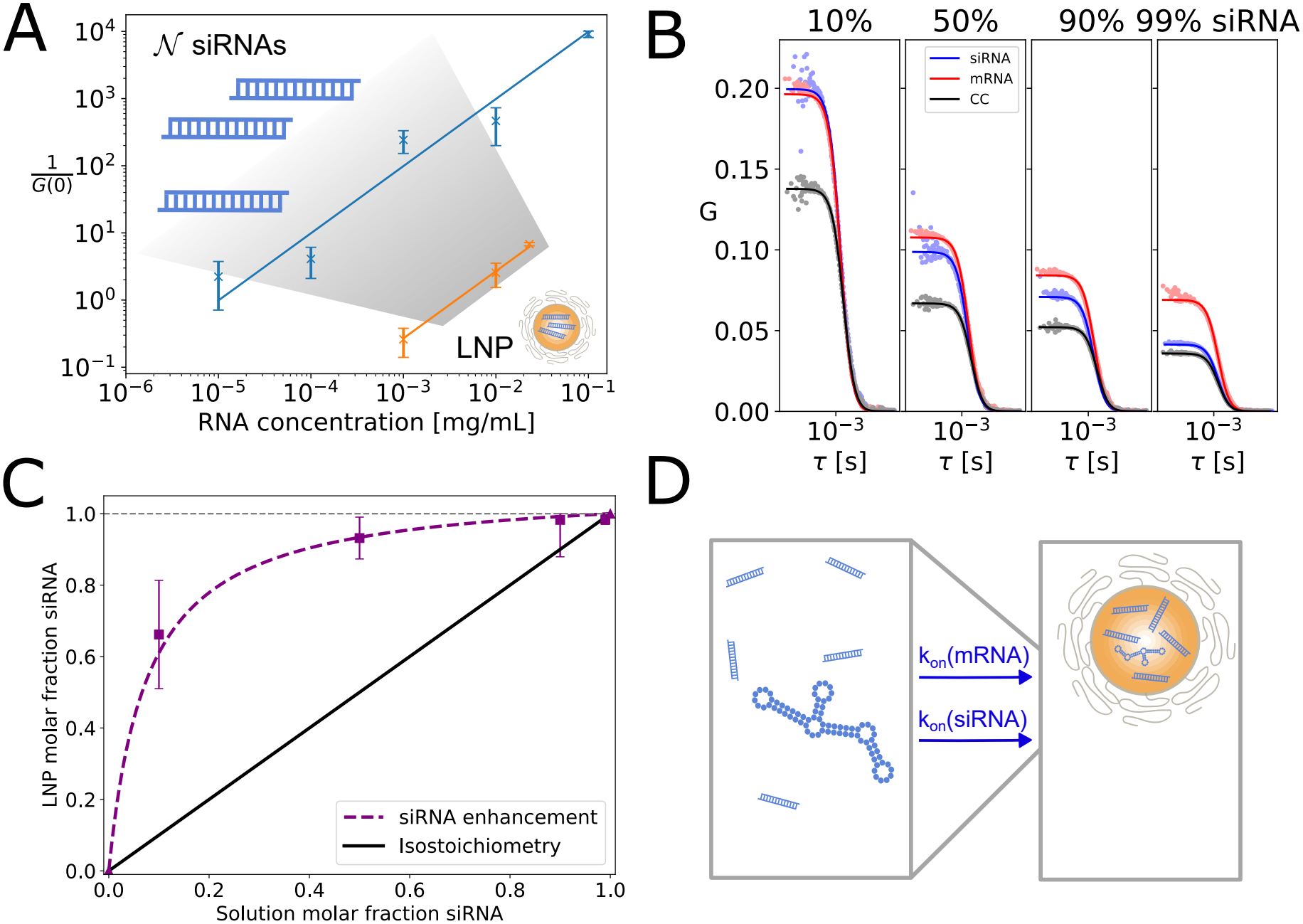
Stoichiometry in siRNA/mRNA coformulations. (A) Dilution series of siRNA (blue) and siRNA-LNPs (orange) yields an siRNA payload of 380 ±50 (*N* = 3). (B) Dual-color FCCS of co-formulated LNPs at 10 % to 99 % molar fraction siRNA. Onset amplitudes are compared to the dilution series to obtain payload of the two species individually. (C) Molar fraction of siRNA inside LNPs (purple) versus molar fraction of siRNA in solution before LNP formation. Note that siRNA content is enhanced compared to the case of unchanged molar ratios (black line). Fixed points of molar fraction at 0 and 1 are given as purple triangles. *N* = 2 replicates. (D) Diffusion-limited model of RNA encapsulation. It is assumed that siRNA and mRNA associate with the LNP forming core at different rates. Hence siRNA encapsulation is enhanced by a factor *α* given by the ratio of on-rates. The resulting molar fraction of siRNA inside LNPs is shown as purple line in (C) (see also main text).

### Stoichiometric payload in siRNA/mRNA-mixed LNPs

In many recent applications the codelivery of both mRNA and siRNA is desired. The ratio between both RNAs can control the expression of both desired and undesirable proteins and it can also reduce production costs.^30^ In this case, LNPs are coformulated by providing an siRNA/mRNA mixture in the microfluidic LNP formation process. As a result both siRNA and mRNA are encapsulated simultaneously and nucleic acid payload inside LNPs is described by two theoretically independent quantities, *N* _siRNA_ and *N* _mRNA_. In the following we define the molar fraction of siRNA in LNPs as

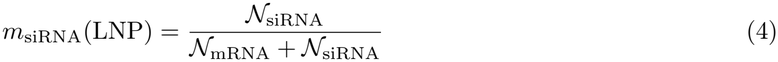

While the molar fraction *m*_siRNA_(0) in the pre-mixing stock solution is precisely known, the resulting molar fraction, *m*_siRNA_(LNP), inside LNPs need not be the same and has to be determined experimentally. In this study, LNPs were loaded with 996 nt Cy5-labeled EGFP mRNA and 20 nt Atto390-labeled siRNA in various molar ratios. Co-formulated LNPs were analyzed by two-color FCS, which determines the auto-correlation function for each fluorescence channel (i.e., the Cy5-mRNA and Atto390-siRNA). Additionally, the cross-correlation function was recorded to report on particles containing both cargos. When probing LNP stoichiometry across a wide range of solution molar fractions *m*_siRNA_(0) = 10 %, 50 %, 90 % and 99 % (Figure 5B), we now obtain two autocorrelation functions for each stoichiometry and the cross-correlation to report on co-diffusion of both. The amplitudes corresponding to mRNA and siRNA in Figure 5B do not necessarily overlap since some particles may contain only one RNA species and the overall ensemble will show fluctuations in payload number. This consequently affects LNP brightness and thus the value of *G*(0). Additionally, we see a decrease in correlation amplitudes with increasing siRNA molar fraction, regardless of the fluorescence channel, suggesting a higher number of LNPs. LNPs with pure mRNA or siRNA are related to molar fractions of *m*_siRNA_(0) = 0 % and 100 %. In mixed cases, we determine each payload individually relying on the dilution series of mRNA and siRNA, where the concentration is reduced by *m*_siRNA_(0).

From both payload numbers, *m*_siRNA_(LNP) is calculated. Mass action kinetics during encapsulation would predict *m*_siRNA_(LNP) = *m*_siRNA_(0), as indicated by a straight line in Figure 5C. However, we observe an enhancement of the molar fraction of siRNA in LNPs (Figure 5C). In order to explain this finding, we propose an aggregation model in which the differences in RNA mobility govern encapsulation efficiency. Within this model siRNA and mRNA encapsulation is limited by kinetic association rates, *k*_on_, during the LNP condensation process (Figure 5D), where the association rates are proportional to the respective diffusion coefficients of the nucleic acid molecules. As siRNA has a higher diffusion coefficient compared to mRNA, more siRNA will arrive in a given complexation time compared to mRNA. To account for this effect, we introduce an siRNA enhancement factor *α*, resulting in a diffusion-limited LNP molar fraction

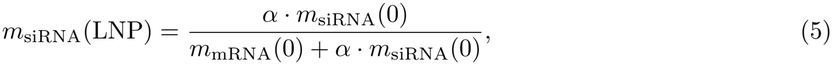

where *m*_mRNA_(0) = 1 −*m*_siRNA_(0). Figure 5C shows a best fit of this expression to the data, yielding *α* = 14.1 ±1.0. In comparison, the diffusion coefficients of siRNA (148 µm^2^ s^*−*1^) and mRNA (13.6 µm^2^ s^*−*1^) as measured by FCS (Figure 3C, Figure 5A) yield a ratio of *D*_siRNA_*/D*_mRNA_ = 10.9 ±1.1, in close agreement with the *α* value derived from the siRNA molar fractions in LNPs. This finding strongly supports the idea that nucleic acid–lipid complexation within LNP formation is governed by the mobility (diffusion coefficient) of the two RNA species.

## Conclusions

In conclusion, we quantified the number of Cy5-labeled mRNA molecules per ONPATTRO-like formulated LNP using TIRF microscopy and fluorescence correlation spectroscopy (FCS). The TIRF analysis used isolated, substrate-immobilized mRNA molecules as a fluorescence intensity-based reference. The FCS analysis determined the mRNA number density from the autocorrelation amplitude and allowed a robust comparison with the number density of LNPs formed upon lipid complexation. Both methods yielded similar payload estimates consistent with values reported in the literature.^21^ When LNP size was varied by adjusting the PEG-lipid content, FCS showed the mRNA payload did not scale with LNP volume, suggesting a nonuni-form RNA distribution within the core and possible association with internal interfaces or substructures, such as blebs. We also show that the particle number of siRNA (380 ±50) is 30 times higher compared to mRNA (12 ±3). This is in agreement with a constant number of base pairs in the LNP assuming charge compensation of the LNP by the nucleic acid backbone. Building on these findings, we determined the stoi-chiometric ratio of siRNA to mRNA in LNPs containing both RNA species. We observed substantial siRNA enrichment relative to the composition of the initial RNA mixture before LNP formation, in agreement with an LNP formation model where nucleic acids exhibit different association rates depending on their diffusion constants. These results show that RNA loading in LNPs is governed by kinetic and structural constraints, underscoring the need for quantitative single-particle studies to guide rational design of multi-component RNA delivery systems.

## Materials and methods

### Microfluidic LNP formulation and characterization

LNPs were prepared by microfluidic mixing using a NanoAssemblr Spark (Cytiva, Freiburg, Germany). Lipids were purchased from Avanti Research (Alabaster, Alabama, USA). DLin-MC3-DMA, DSPC, cholesterol and DMG-PEG2000 were dissolved in ethanol and combined at a molar ratio of 50:10:38.5:1.5, following the ONPATTRO composition. The organic phase was mixed with an aqueous phase containing mRNA in 50 mM citrate buffer (pH 3) at a volume ratio of 1:2 (organic:aqueous). Mixing conditions were adjusted to yield a final mRNA concentration of 50 µg mL^*−*1^ and an N/P ratio of 3. Enhanced green fluorescent protein (eGFP) mRNA (996 nt) was obtained from TriLink BioTechnologies (San Diego, CA, USA). The mRNA was labeled with Cy5 via partial substitution of uridine residues (25 % Cyanine5-UTP, 75 % 5-methoxy-UTP), corresponding to an approximate labeling efficiency of 25 % of uridine positions, as specified by the manufacturer. The transcript was capped (Cap1) using CleanCap technology. The siRNA sequence used was 5’-GCA AGC UGA CCC UGA AGU UC-3’, labeled with Atto390 (Kaneka Eurogentec, Seraing, Belgium). Formulated LNPs were dialyzed using Slide-A-Lyzer MINI dialysis devices (3.5 kDa molecular weight cut-off; Thermo Fisher Scientific, Waltham, MA, USA) against Milli-Q water for 12 h at room temperature to remove ethanol and exchange buffer. Centrifugation at 20 000 rcf for 1 h effectively reduced the presence of large aggregates, visible as intensity bursts in the recorded trace for FCS. Removal of these aggregates was necessary to enable subsequent fluorescence correlation spectroscopy measurements. Particle size and polydispersity index were determined by DLS (DynaPro NanoStar, Wyatt Technology). Encapsulation efficiency was measured using the Quant-iT RiboGreen assay (Invitrogen) using 2 % Triton X-100 in water as detergent. mRNA concentration was determined by UV absorbance using a NanoDrop spectrophotometer.

### Variation of LNP size

Under “PEG crowding”, PEG lipid (molar fraction *x*) is added at the expense of cholesterol (MC3:DSPC:Chol:PEG = 50:10:40 −*x*:*x*). Our “fixed surface composition” approach keeps the ratios of molar fractions of bulk lipids MC3/cholesterol and surface lipids PEG/DSPC constant at the ONPATTRO reference values, but varies the proportion of bulk to surface lipids PEG/MC3: (MC3:DSPC:Chol:PEG) = (*ax, b*(100− *x*), (1 −*a*)*x*, (1 −*b*)(100 −*x*)), where *a* = MC3*/*(MC3+Chol) = const. and *b* = DSPC*/*(DSPC + PEG) = const.

### TIRF-based quantification of mRNA loading in LNPs with spectral correction

Glass slides (Ibidi, Martinsried, Germany, Cat. No. 80828) were functionalized by oxygen plasma treatment, followed by incubation with 1 % BSA in PBS. After washing, slides were incubated with neutravidin and washed again with PBS. PEG-biotinylated LNPs were immobilized on the functionalized surface via biotin–neutravidin in-teractions by incubation at an mRNA concentration of 1 µg mL^*−*1^. Naked Cy5-labeled mRNA was hybridized to biotinylated polyT strands and tethered to the surface using the same linkage chemistry. Imaging was performed on a Nikon Ti Eclipse microscope using TIRF illumination at 640 nm with an exposure time of 800 ms and a laser power of 0.3 mW. A CFI Apochromat TIRF 60× oil objective and a 432/515/595/730 HC quadband filter were used. Imaging conditions were selected to balance signal intensity and photobleaching (bleaching half-time *t*_1*/*2_ = 10 s). To ensure even illumination, only the center region of the image was used. Localization and quantification of fluorescence spots were performed using the ThunderSTORM plugin for ImageJ.^31^ For each detected particle, the integrated intensity and apparent size were extracted. Photon count histograms for LNPs were fitted using a Poisson distribution *p*(*k*) = *µ*^*k*^*e*^*−µ*^*/k*!. The expectation value ⟨*I*(LNP) ⟩ was obtained as fit parameter. Count histograms of mRNA were fitted with an exponential distribution. A Gaussian kernel density estimate was used to obtain *I*(mRNA = 1). Standard error of the mean and fit uncertainties were used to estimate errors.

Emission spectra of encapsulated and free Cy5-mRNA before and after release induced by 2 % Triton X-100 were recorded using a JASCO FP-8550 fluorescence spectrometer. To account for differences in detection efficiency due to the microscope filter set, a correction factor *ψ*_T_ was introduced:

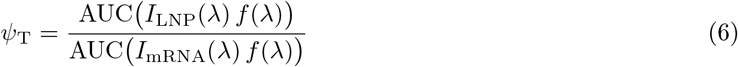

where *f* (*λ*) is the transmission function of the HC quadband filter and AUC gives the integrated area under the curve over the detection window (600 nm to 800 nm). A value of *ψ*_T_ = 1.27 was obtained.

### Fluorescence correlation spectroscopy

FCS measurements were performed on a ZEISS LSM980 confocal microscope using a 40× water immersion objective (LD LCI Plan-Apochromat 40x/1.2 Imm Corr DIC M27, 420862-9970-000) and an Ibidi 6-channel slide (Cat. No. 80608) at 639 nm (Cy5-mRNA) and 405 nm (Atto390-siRNA) laser lines. Confocal volumes were calibrated using standards of known diffusion coefficients: 250 µm^2^ s^*−*1^ (Cy5)^32^ and 440 µm^2^ s^*−*1^ (Atto390).^33^ All confocal waists were found to be between 200 and 350 nm and confocal volumes between 0.2 and 1 fL. Each independent sample was measured with 100 repeats of 10 s each. For each composition 3 independently formulated samples were measured. Autocor-relation curves were fitted with a one-component 3D normal diffusion model with triplet component. Size measurements of hydrodynamic radius are given via the Stokes–Einstein relation. Because the LNP radius amounts to approximately 20 % of the lateral confocal width *w*_*r*_, a finite-size correction for diffusion times and particle number was applied.^34^

### Determination of siRNA/mRNA stoichiometry using two-color FCS

Measurements were taken on a ZEISS LSM780 confocal microscope (objective: LD LCI Plan-Apochromat 40x/1.2 Imm Corr DIC M27, 420862-9970-000). The setup was prepared as described in a previous protocol for cross-correlation spectroscopy.^35^ LNP co-formulations were produced at constant N/P = 3 to adjust for the double charge density on the double-stranded siRNA and to keep the electrostatic environment equivalent. Autocorrelation curves were fitted with a 3D normal diffusion model with triplet component to obtain the onset amplitude.

### Cryo-EM imaging of LNPs

A 3 µL aliquot of freshly prepared LNPs was applied to a glow-discharged holey gold coated grid (300 mesh Au-flat 1.2/1.3), blotted with filter paper and plunge-frozen in liquid ethane using an Automatic Plunge Freezer EM GP2 (Leica Microsystems). The data were recorded on a JEM-2100F transmission electron microscope (JEOL) with a mounted F416 CCD camera (TVIPS). Measurements of LNP morphology were carried out by manual segmentation of well-separated LNPs based on their internal structure, followed by characterization of the segmentation masks in Fiji.^36^

## Supporting information

The Supporting Information is available as a separate file accompanying this preprint.

- Supporting Information (PDF): materials and methods for spectroscopy; excitation and emission spectra as well as fluorescence lifetime data of bare and encapsulated mRNA; DLS measurements; Cryo-EM size distributions

## Author information

### Present addresses

Thomas Kellerer: MPI for Biochemistry, 82152 Planegg, Germany

### Author contributions

B.K., J.H., T.K. and J.R. conceived the study. B.K. designed and performed experiments, analyzed and interpreted data. D.M. designed and performed fluorescence spectroscopy and lifetime experiments and analyzed and interpreted data with T.O.P. and T.C. N.S. performed the Cryo-EM imaging. He designed these experiments and analyzed the Cryo-EM data together with M.F. B.K. and D.M. wrote the manuscript, which was edited by T.C. and J.R. All authors have given approval to the final version of the manuscript.

### Funding sources

This work was supported by the German Research Foundation (RA 655/12-1, CO 879/6-1), by the Free State of Bavaria under the Excellence Strategy through the ONE MUNICH Project Munich Multiscale Biofabrication, TU Dortmund University and through the Federal Ministry of Research, Technology and Space (BMFTR), Germany under Project 05K2026 - 2025-08197 LIPION within the frame-work of the Swedish-German research collaboration Röntgen-Ångström Cluster.

### Notes

T.C. declares commercial interest in FluoroBrick Solutions GmbH, a company that distributes micro-spectroscopy setups. All other authors declare no competing financial interest.

## Acknowledgements

B.K., J.H. and J.R. thank Eric Olsen and Sabrina Leslie for fruitful discussion. The technical and scientific support provided by Gerlinde Schwake is gratefully acknowledged. B.K. thanks Jan-Hagen Krohn for useful discussion and advice.

## Abbreviations

LNP: lipid nanoparticle
FCS: fluorescence correlation spectroscopy
TIRF: total internal reflection fluorescence

## Notes

### Summary of Updates

Erroneous axis unit in FCS autocorrelation plots corrected from ms to s.

## References

(1) Cullis, P. R.; Hope, M. J. Lipid Nanoparticle Systems for Enabling Gene Therapies. Mol. Ther. 2017,25, 1467–1475.

(2) Adams, D.; Gonzalez-Duarte, A.; O’Riordan, W. D.; Yang, C.-C.; Ueda, M.; Kristen, A. V.; Tournev, I.; Schmidt, H. H.; Coelho, T.; Berk, J. L.; Lin, K.-P.; Vita, G.; Attarian, S.; Planté-Bordeneuve, V.; Mezei, M. M.; Campistol, J. M.; Buades, J.; Brannagan, T. H.; Kim, B. J.; Oh, J.; Parman, Y.; Sekijima, Y.; Hawkins, P. N.; Solomon, S. D.; Polydefkis, M.; Dyck, P. J.; Gandhi, P. J.; Goyal, S.; Chen, J.; Strahs, A. L.; Nochur, S. V.; Sweetser, M. T.; Garg, P. P.; Vaishnaw, A. K.; Gollob, J. A.;Suhr O. B. Patisiran, an RNAi Therapeutic, for Hereditary Transthyretin Amyloidosis. N. Engl. J. Med. 2018, 379, 11–21.

(3) Brimacombe, C. A.; Kulkarni, J. A.; Cheng, M. H.; An, K.; Witzigmann, D.; Cullis, P. R. Rational design of lipid nanoparticles for enabling gene therapies. Mol. Ther. Methods Clin. Dev. 2025, 33, 101518.

(4) Xue, L.; Ang, M. J. Y.; Swingle, K. L.; Fitzgerald, E.; Chalom, O.; Gu, N.; Mitchell, M. J. Structural evolution of ionizable lipids for nucleic acid delivery. Nat. Rev. Chem. 2026, DOI: 10.1038/s41570-026-00865-0.

(5) Yanez Arteta, M.; Kjellman, T.; Bartesaghi, S.; Wallin, S.; Wu, X.; Kvist, A. J.; Dabkowska, A.; Székely, N.; Radulescu, A.; Bergenholtz, J.; Lindfors, L. Successful reprogramming of cellular protein production through mRNA delivered by functionalized lipid nanoparticles. Proc. Natl. Acad. Sci. U.S. A. 2018, 115, DOI: 10.1073/pnas.1720542115.

(6) Uebbing, L.; Ziller, A.; Siewert, C.; Schroer, M. A.; Blanchet, C. E.; Svergun, D. I.; Ramishetti, S.; Peer, D.; Sahin, U.; Haas, H.; Langguth, P. Investigation of pH-Responsiveness inside Lipid Nanoparticles for Parenteral mRNA Application Using Small-Angle X-ray Scattering. Langmuir 2020, 36, 13331–13341.

(7) Philipp, J.; Dabkowska, A.; Reiser, A.; Frank, K.; Krzysztoń, R.; Brummer, C.; Nickel, B.; Blanchet,C. E.; Sudarsan, A.; Ibrahim, M.; Johansson, S.; Skantze, P.; Skantze, U.; Östman, S.; Johansson, M.; Henderson, N.; Elvevold, K.; Smedsrød, B.; Schwierz, N.; Lindfors, L.; Rädler, J. O. pH-dependent structural transitions in cationic ionizable lipid mesophases are critical for lipid nanoparticle function. Proc. Natl. Acad. Sci. U. S. A. 2023, 120, e2310491120.

(8) Yu, H.; Angelova, A.; Angelov, B.; Dyett, B.; Matthews, L.; Zhang, Y.; El Mohamad, M.; Cai, X.; Valimehr, S.; Drummond, C. J.; Zhai, J. Real-Time pH-Dependent Self-Assembly of Ionisable Lipids from COVID-19 Vaccines and In Situ Nucleic Acid Complexation. Angew. Chem., Int. Ed. 2023, 62, e202304977.

(9) Zhang, Y.; Leung, J.; Cheng, J.; Momeni, A.; Kamanzi, A.; Bido, A. T.; Liao, S.; Kang, E.; Strong, C.; Kastrup, C. J.; Leslie, S.; Cheng, M. H. Y.; Cullis, P. R. Increasing Lipid Nanoparticle mRNA Copy Number Improves Transfection Potency. Adv. Mater. 2026, e75174.

(10) Müller, J. A.; Schäffler, N.; Kellerer, T.; Schwake, G.; Ligon, T. S.; Rädler, J. O. Kinetics of RNA-LNP delivery and protein expression. Eur. J. Pharm. Biopharm. 2024, 197, 114222.

(11) Sebastiani, F.; Yanez Arteta, M.; Lerche, M.; Porcar, L.; Lang, C.; Bragg, R. A.; Elmore, C. S.; Krishnamurthy, V. R.; Russell, R. A.; Darwish, T.; Pichler, H.; Waldie, S.; Moulin, M.; Haertlein, M.; Forsyth, V. T.; Lindfors, L.; Cárdenas, M. Apolipoprotein E Binding Drives Structural and Compositional Rearrangement of mRNA-Containing Lipid Nanoparticles. ACS Nano 2021, 15, 6709–6722.

(12) Cui, L.; Pereira, S.; Sonzini, S.; van Pelt, S.; Romanelli, S. M.; Liang, L.; Ulkoski, D.; Krishnamurthy,V. R.; Brannigan, E.; Brankin, C.; Desai, A. S. Development of a high-throughput platform for screening lipid nanoparticles for mRNA delivery. Nanoscale 2022, 14, 1480–1491.

(13) Nawaz, M.; Heydarkhan-Hagvall, S.; Tangruksa, B.; González-King Garibotti, H.; Jing, Y.; Maugeri, M.; Kohl, F.; Hultin, L.; Reyahi, A.; Camponeschi, A.; Kull, B.; Christoffersson, J.; Grimsholm, O.; Jennbacken, K.; Sundqvist, M.; Wiseman, J.; Bidar, A. W.; Lindfors, L.; Synnergren, J.; Valadi, H.Lipid Nanoparticles Deliver the Therapeutic VEGFA mRNA In Vitro and In Vivo and Transform Extracellular Vesicles for Their Functional Extensions. Adv. Sci. 2023, 10, 2206187.

(14) Carrasco, M. J.; Alishetty, S.; Alameh, M.-G.; Said, H.; Wright, L.; Paige, M.; Soliman, O.; Weissman, D.; Cleveland, T. E.; Grishaev, A.; Buschmann, M. D. Ionization and structural properties of mRNA lipid nanoparticles influence expression in intramuscular and intravascular administration. Commun.Biol. 2021, 4, 956.

(15) Sabnis, S.; Kumarasinghe, E. S.; Salerno, T.; Mihai, C.; Ketova, T.; Senn, J. J.; Lynn, A.; Bulychev, A.; McFadyen, I.; Chan, J.; Almarsson, O.; Stanton, M. G.; Benenato, K. E. A novel amino lipid series for mRNA delivery: Improved endosomal escape and sustained pharmacology and safety in non-human primates. Mol. Ther. 2018, 26, 1509–1519.

(16) Kamanzi, A.; Gu, Y.; Tahvildari, R.; Friedenberger, Z.; Zhu, X.; Berti, R.; Kurylowicz, M.; Witzig-mann, D.; Kulkarni, J. A.; Leung, J.; Andersson, J.; Dahlin, A.; Höök, F.; Sutton, M.; Cullis, P. R.; Leslie S. Simultaneous, Single-Particle Measurements of Size and Loading Give Insights into the Structure of Drug-Delivery Nanoparticles. ACS Nano 2021, 15, 19244–19255.

(17) Graves, S.; Jasinski, M.; Olsén, E.; Kamanzi, A.; Zhang, Y.; Leung, J.; Venier-Karzis, M.; Safaeesirat, A.; Cullis, P. R.; Leslie, S. R. Direct Counting of mRNA Copies Inside Individual Lipid Nanoparticles Using In Situ Lysis and Labeling. Anal. Chem. 2026, 98, 15961–15967.

(18) Li, S.; Hu, Y.; Li, A.; Lin, J.; Hsieh, K.; Schneiderman, Z.; Zhang, P.; Zhu, Y.; Qiu, C.; Kokkoli, E.; Wang, T.-H.; Mao, H.-Q. Payload distribution and capacity of mRNA lipid nanoparticles. Nat. Commun. 2022, 13, 5561.

(19) Li, S.; Hu, Y.; Lin, J.; Schneiderman, Z.; Shao, F.; Wei, L.; Li, A.; Hsieh, K.; Kokkoli, E.; Curk, T.; Mao, H.-Q.; Wang, T.-H. Single-Particle Spectroscopic Chromatography Reveals Heterogeneous RNA Loading and Size Correlations in Lipid Nanoparticles. ACS Nano 2024, 18, 15729–15743.

(20) Sanchez-Fernandez, A.; Donnelly, K. A.; Bolinsson, H.; Börjesdotter, A.-M.; Rønnemoes Bobak, T.; Erlendsson, S.; Mohammadi-Zerankeshi, M.; AboulFotouh, K.; Kawelah, M. R.; Herranz-Trillo, F.; Schagerlöf, H.; Capasso Palmiero, U.; Huus, K.; Johnston, K. P.; Marras, A. E.; Nilsson, L. Structural heterogeneity in mRNA-LNP subpopulations revealed by AF4-SAXS: implications for cargo loadingand cell transfection. bioRxiv 2026, 2026.01.16.699683.

(21) Olsén, E.; Kirchmair, B.; Rädler, J. O.; Leslie, S. R. Normalized Nucleic Acid Payload in Lipid Nanoparticles: A Perspective on the Harmonization of Published Data. ChemRxiv 2026, DOI: 10.26434/chemrxiv.15011677/v1.

(22) Kamanzi, A.; Tuckmantel Bido, A.; Zhang, Y.; Olsén, E.; Stibbards-Lyle, M.; Jasinski, M.; Gu, Y.; Wang, B.; Venier-Karzis, M.; Berti, R.; Jeliazkova, M.; Shaheen, C.; Leung, J.; Cheng, M. H. Y.; Cullis,P. R.; Leslie, S. Single-Particle Multiparametric Microscopy Reveals Structural, Size, and Payload Heterogeneity in mRNA-Loaded Lipid Nanoparticles. ACS Nano 2026, 20, 1290–1303.

(23) Levitus, M.; Ranjit, S. Cyanine dyes in biophysical research: the photophysics of polymethine fluorescent dyes in biomolecular environments. Q. Rev. Biophys. 2011, 44, 123–151.

(24) Gebhardt, C.; Lehmann, M.; Reif, M. M.; Zacharias, M.; Gemmecker, G.; Cordes, T. Molecular and Spectroscopic Characterization of Green and Red Cyanine Fluorophores from the Alexa Fluor and AF Series. ChemPhysChem 2021, 22, 1566–1583.

(25) Schwille, P.; Haustein, E. Fluorescence Correlation Spectroscopy: An Introduction to its Concepts and Applications. In Biophysics Textbook Online; Biophysical Society: 2001.

(26) Krichevsky, O.; Bonnet, G. Fluorescence correlation spectroscopy: the technique and its applications.Rep. Prog. Phys. 2002, 65, 251.

(27) Hassett, K. J.; Higgins, J.; Woods, A.; Levy, B.; Xia, Y.; Hsiao, C. J.; Acosta, E.; Almarsson, Ö.; Moore, M. J.; Brito, L. A. Impact of lipid nanoparticle size on mRNA vaccine immunogenicity. J. Controlled Release 2021, 335, 237–246.

(28) Biru, K. E.; Fiedler, B. M.; Gurnani, P. Mushrooms to brushes: Tackling the PEG dilemma in mRNA lipid nanoparticles with complex polymers. Matter 2025, 8, DOI: 10.1016/j.matt.2025.102386.

(29) Hammel, M.; Fan, Y.; Sarode, A.; Byrnes, A. E.; Zang, N.; Kou, P.; Nagapudi, K.; Leung, D.; Hoogenraad, C. C.; Chen, T.; Yen, C.-W.; Hura, G. L. Correlating the Structure and Gene Silencing Activity of Oligonucleotide-Loaded Lipid Nanoparticles Using Small-Angle X-ray Scattering. ACS Nano 2023,17, 11454–11465.

(30) Ball, R. L.; Hajj, K. A.; Vizelman, J.; Bajaj, P.; Whitehead, K. A. Lipid Nanoparticle Formulations for Enhanced Co-delivery of siRNA and mRNA. Nano Lett. 2018, 18, 3814–3822.

(31) Ovesný, M.; Křížek, P.; Borkovec, J.; Švindrych, Z.; Hagen, G. M. ThunderSTORM: a comprehensive ImageJ plug-in for PALM and STORM data analysis and super-resolution imaging. Bioinformatics 2014, 30, 2389–2390.

(32) Widengren, J.; Schwille, P. Characterization of Photoinduced Isomerization and Back-Isomerization of the Cyanine Dye Cy5 by Fluorescence Correlation Spectroscopy. J. Phys. Chem. A 2000, 104, 6416–6428.

(33) Van Zanten, C.; Ryder, A. G. Estimating Poly(N-isopropylacrylamide) size in solution below the LCST using fluorescence correlation spectroscopy with non-covalent bound fluorophores. Polym. Test. 2024, 137, 108500.

(34) Wu, B.; Chen, Y.; Müller, J. D. Fluorescence Correlation Spectroscopy of Finite-Sized Particles. Biophys. J. 2008, 94, 2800–2808.

(35) Bacia, K.; Schwille, P. Practical guidelines for dual-color fluorescence cross-correlation spectroscopy.Nat. Protoc. 2007, 2, 2842–2856.

(36) Schindelin, J.; Arganda-Carreras, I.; Frise, E.; Kaynig, V.; Longair, M.; Pietzsch, T.; Preibisch, S.; Rueden, C.; Saalfeld, S.; Schmid, B.; Tinevez, J.-Y.; White, D. J.; Hartenstein, V.; Eliceiri, K.; Tomancak, P.; Cardona, A. Fiji: an open-source platform for biological-image analysis. Nat. Methods 2012, 9, 676–682.

