## Supplementary material for "siRNA/mRNA payload of lipid nanoparticles quantified using fluorescence microscopy and correlation spectroscopy": SI

#### Materials and methods used for data in the Supporting Information

**UV–Vis spectroscopy.** Fluorescence spectra were recorded using a JASCO FP-8550 fluorescence spectrometer and a Hellma® high-precision quartz ultra-micro cuvette (3 mm × 3 mm path length, 15 mm center height, dimensions 45 mm × 12.5 mm × 12.5 mm). The measurements were carried out as follows: Freshly prepared lipid nanoparticles containing mRNA-Cy5 at a concentration of $4\,\mathrm{mg\,L^{-1}}$ were first measured in the cuvette. For measurements involving Triton™ X-100 (TX), a TX stock solution in water (2 %) was added directly to the same cuvette at a 1:1 volume ratio. To account for the resulting dilution, the spectral intensities of TX-treated samples were multiplied by a factor of 2 to match the mRNA-Cy5 concentration of the untreated samples. For emission and excitation spectra the experimental parameters were $\lambda_{\mathrm{ex}} = 595\,\mathrm{nm}$ and $\lambda_{\mathrm{em}} = 730\,\mathrm{nm}$, respectively, and $\Delta\lambda_{\mathrm{ex}} = \Delta\lambda_{\mathrm{em}} = 5\,\mathrm{nm}$.

**Fluorescence decay recording and analysis.** Fluorescence decays were obtained by time-correlated single-photon counting (TCSPC). Experiments were performed on an inverted microscope with a 60× water immersion objective (Nikon M Plan Apo, NA 1.20) with time-resolved single photon detection (MicroTime 200, PicoQuant) featuring single photon counting electronics with picosecond time resolution (HydraHarp 400, PicoQuant). Excitation was achieved using a linearly polarized pulsed laser diode (LDH-D-C-640, 636 nm, PicoQuant) operating at a repetition period of 25 ns and a power of 10 µW at the objective. Emitted photons were registered by single-photon avalanche diodes (SPCM-AQR-14, PerkinElmer) after passing through a red band-pass filter (ET700/75M, Chroma).

The registered photon arrival times were binned into fluorescence decay histograms using the software ChiSurf[1] at a temporal resolution of 32 ps per bin. The resulting fluorescence decay curves, $f(t)$, were fitted by multi-exponential models,

$$f(t) = \sum_{i=1}^{N} x^{(i)} \cdot e^{-t/\tau^{(i)}}, \tag{S1}$$

where $x^{(i)}$ and $\tau^{(i)}$ denote the species fractions and fluorescence lifetimes, respectively, and $N$ is the number of fluorescence decay components. For multi-exponential models the fluorescence intensity weighted average lifetime $\langle\tau\rangle_{\mathrm{Int}}$ was computed as

$$\langle\tau\rangle_{\mathrm{Int}} = \sum_{i=1}^{N} x^{(i)} \cdot \left(\tau^{(i)}\right)^2 . \tag{S2}$$

The model parameters were obtained by iterative reconvolution fitting, incorporating the experimentally measured instrument response function (IRF). The IRF was determined from scattered light in water under the same detection conditions as used for the sample. During parameter optimization, the experimental nuisance parameters were simultaneously optimized, including a constant non-fluorescent background contribution (e.g., detector dark counts), a temporal offset between the IRF and the fluorescence decay, and remaining scattered light contributions. The optimization aimed to minimize the reduced chi-square statistic, $\chi^2_{\mathrm{r}}$, while goodness of fit was additionally evaluated by examining the weighted residuals.

### Supplementary data

#### Fluorescence spectra and lifetimes

The intensity-averaged lifetimes are summarized in Table S1; spectra and decays are shown in Figure S1.

Table S1: Intensity-averaged fluorescence lifetimes. The intact LNP has an increased lifetime compared to naked mRNA and the Triton-treated LNP.

| | LNP mRNA-Cy5 | LNP mRNA-Cy5 + 1% Triton X | mRNA-Cy5 | mRNA-Cy5 + 1% Triton X |
|---|---|---|---|---|
| $\langle\tau\rangle_{\mathrm{Int}}$ [ns] | $1.98 \pm 0.05$ | $1.58 \pm 0.03$ | $1.49 \pm 0.02$ | $1.54 \pm 0.05$ |
| Independent repeats | 3 | 2 | 3 | 2 |

#### DLS measurements

DLS data are shown in Figure S2.

#### Cryo-TEM size distributions of LNPs

Cryo-TEM size distributions are shown in Figure S3.

### References

(1) Peulen, T.-O. Exploring Time-Resolved Fluorescence Data: A Software Solution for Model Generation and Analysis. *Spectrosc. J.* **2025**, *3*, 16.

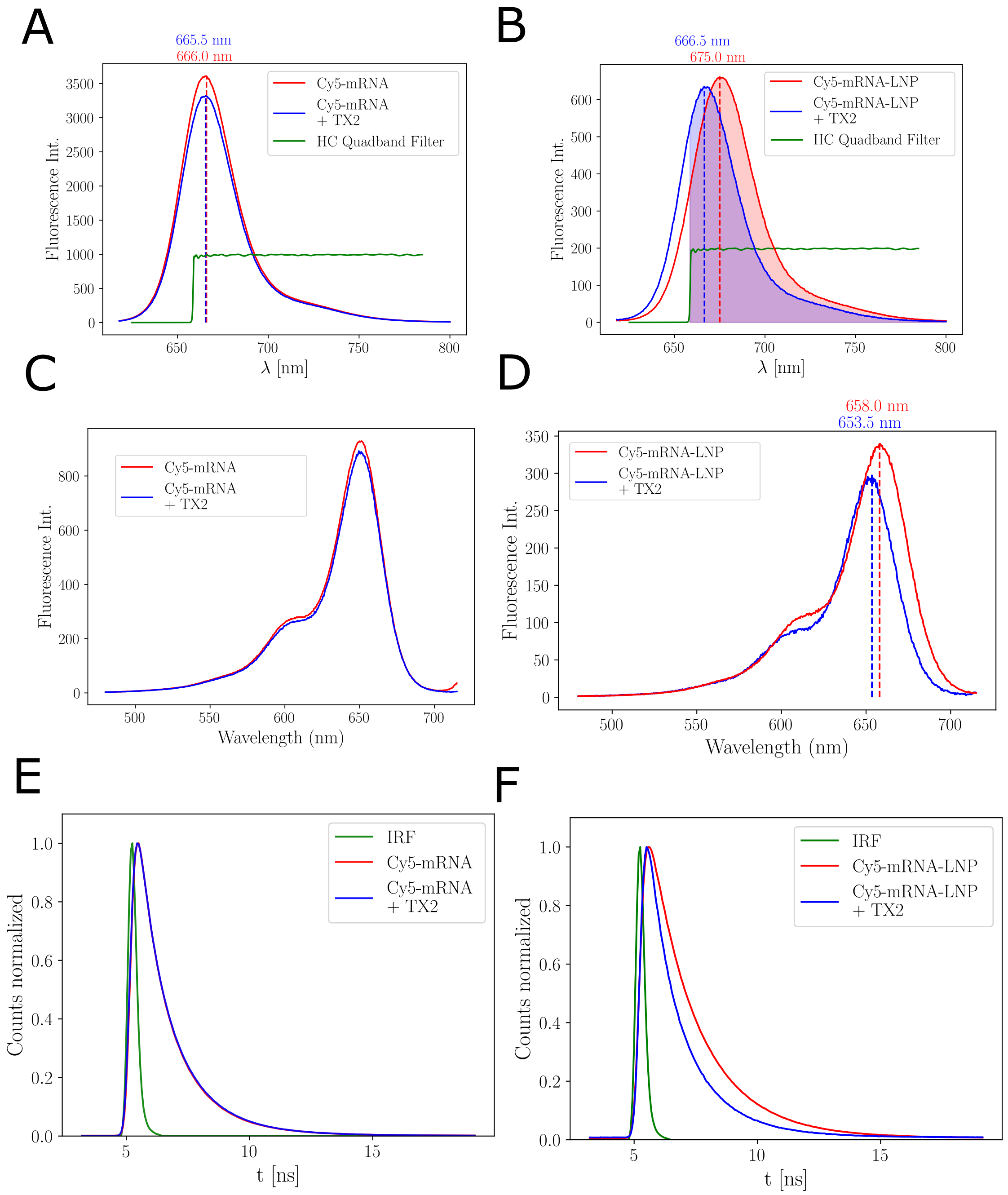


Figure S1: Emission spectra of Cy5-mRNA (A) and Cy5-mRNA-LNPs (B) at an excitation wavelength of 595 nm. Between the water solution (red) and treatment with TX for LNP disruption (blue) a hypsochromic shift of $(8.5 \pm 0.5)$ nm in the peak position occurs. Given by the used HC Quadband filter FF01-432/515/595/730-25 (green), a brightness correction factor of $\psi_\mathrm{T} = 1.27$ between the shaded areas is obtained. The peaks in the excitation spectra of Cy5-mRNA (C) and Cy5-mRNA-LNPs (D) are also shifted by $(4.5 \pm 0.5)$ nm before treatment with TX (red) and after (blue). Fluorescence lifetime decays of (E) Cy5-mRNA and (F) Cy5-mRNA-LNPs in water (red) and treated with 1 % Triton X (blue) at an excitation wavelength of 636 nm. The intact LNP has an increased lifetime compared to naked mRNA and the Triton-treated LNP.

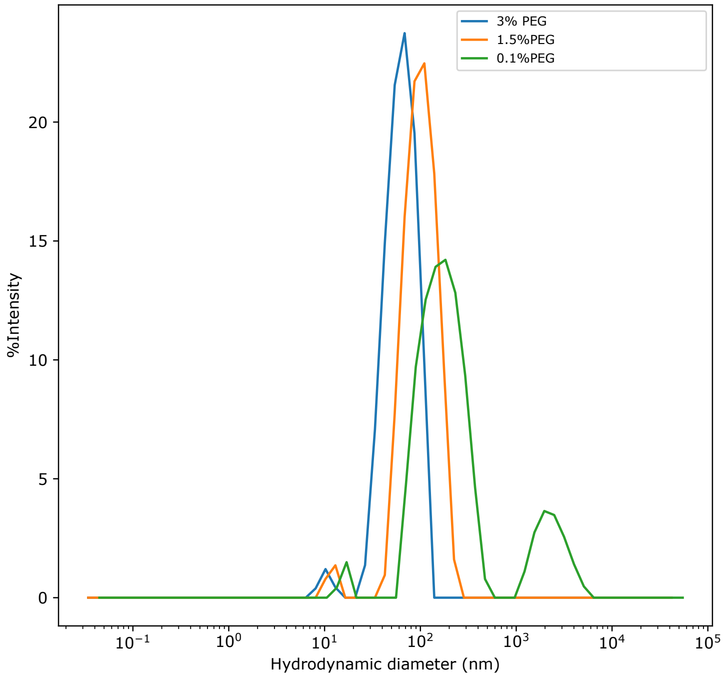


Figure S2: Intensity-weighted DLS size distributions of LNPs with 3 mol % (blue), 1.5 mol % (orange) and 0.1 mol % (green) PEG-lipid. At 0.1 mol % PEG-lipid the main peak broadens and an additional population in the micrometre range appears, indicating aggregation upon reduced LNP stability.

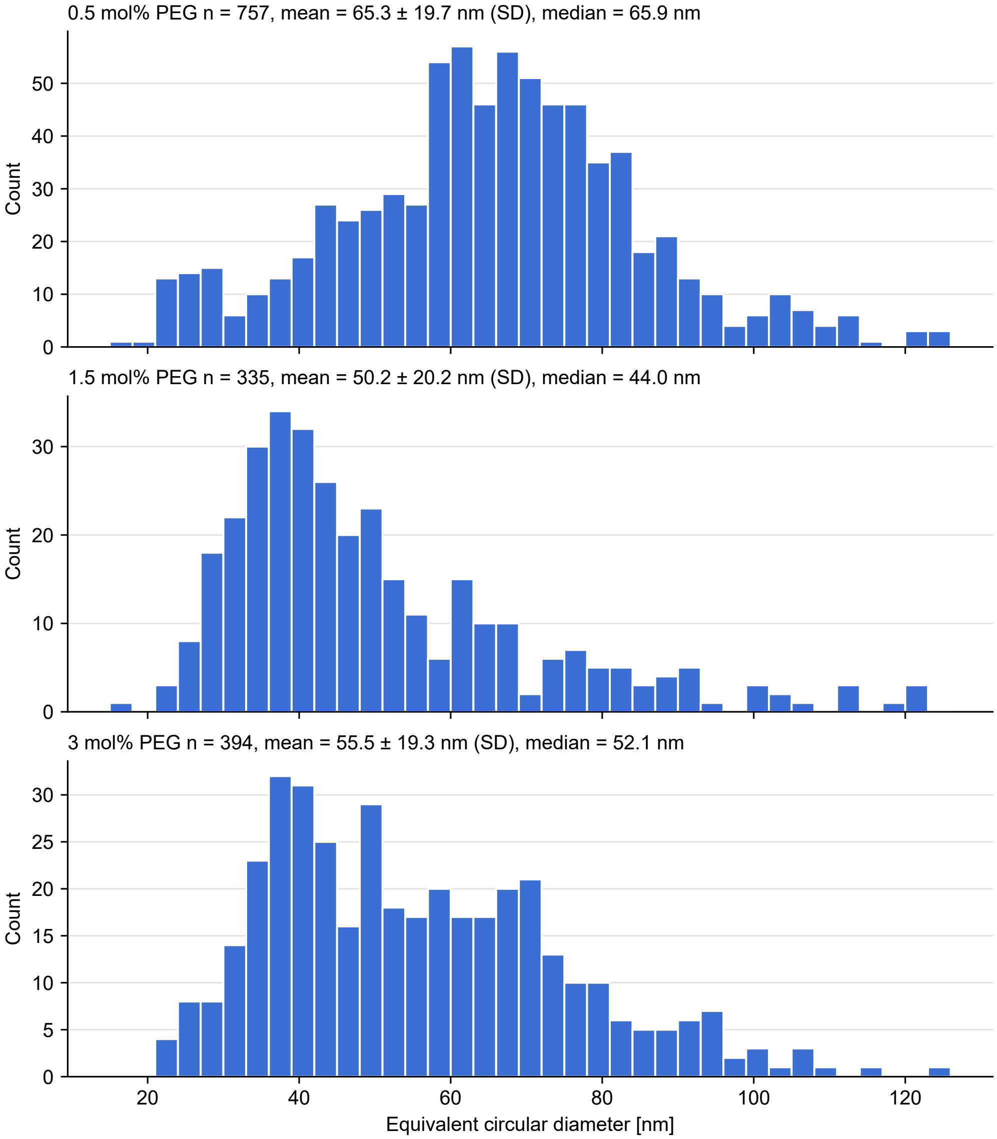


Figure S3: Cryo-TEM size distributions of LNPs with 0.5 mol %, 1.5 mol % and 3 mol % DMG-PEG2k. Histograms of the equivalent circular diameter, $d = \sqrt{4A/\pi}$, where $A$ is the projected area of each segmented particle in Fiji. Only particles with $15\,\mathrm{nm} \leq d \leq 125\,\mathrm{nm}$, circularity $\geq 0.7$ and aspect ratio $\leq 1.5$ were included. Bin width is 3 nm. Number of particles ($n$), mean ± SD and median are given above each panel.
